# A sample preparation procedure enables acquisition of 2-channel super-resolution 3D STED image of an entire oocyte

**DOI:** 10.1101/2023.03.07.531472

**Authors:** Michaela Frolikova, Michaela Blazikova, Martin Capek, Helena Chmelova, Jan Valecka, Veronika Kolackova, Eliska Valaskova, Martin Gregor, Katerina Komrskova, Ondrej Horvath, Ivan Novotny

**Affiliations:** Laboratory of Reproductive Biology, Institute of Biotechnology of the Czech Academy of Sciences, BIOCEV, Prumyslova 595. 252 50 Vestec, Czech Republic; Light Microscopy Core Facility, Institute of Molecular Genetics of the Czech Academy of Sciences, Videnska 1083, 142 20 Prague 4, Czech Republic; Laboratory of Biomathematics, Institute of Physiology of the Czech Academy of Sciences, Videnska 1083, 142 20 Prague 4, Czech Republic; Centre of the Region Hana for Biotechnological and Agricultural Research, Institute of Experimental Botany of the Czech Academy of Sciences, Slechtitelu 31, 779 00 Olomouc, Czech Republic; Laboratory of Integrative Biology, Institute of Molecular Genetics of the Czech Academy of Sciences, Videnska 1083, 142 20 Prague 4, Czech Republic; Department of Zoology, Faculty of Science, Charles University, Vinicna 7, 128 44 Prague 2, Czech Republic

**Keywords:** Sample preparation, super-resolution, 3D STED, oocyte

## Abstract

Super-resolution (SR) microscopy is a cutting-edge method that can provide detailed structural information with high resolution. However, the thickness of the specimen has been a major limitation for SR methods, and larger structures have posed a challenge. To overcome this, the key step is to optimize sample preparation to ensure optical homogeneity and clarity, which can enhance the capabilities of SR methods for the acquisition of thicker structures.

Oocytes are the largest cells in the mammalian body and are crucial objects in reproductive biology. They are especially useful for studying membrane proteins. However, oocytes are extremely fragile and sensitive to mechanical manipulation and osmotic shocks, making sample preparation a critical and challenging step.

We present an innovative, simple, and sensitive approach to oocyte sample preparation for 3D STED acquisition. This involves alcohol dehydration and mounting into a high refractive index medium. This extended preparation procedure allowed us to successfully obtain a unique 2-channel 3D STED super-resolution image of an entire mouse oocyte.

By optimizing sample preparation, we can overcome the limitations of SR methods and obtain high-resolution images of larger structures, such as oocytes, Knowledge of which are important for understanding fundamental biological processes.

**RESEARCH HIGHLIGHTS:**

- This study aimed to develop a successful sample preparation protocol for imaging mouse oocytes using 3D STED super-resolution microscopy.
- The results showed that the oocyte sample was optically homogenous, enhancing the capability of the 3D STED method to capture high-resolution images throughout the full depth of the sample, resulting in highly similar SR images.
- The 3D STED point spread function (PSF) pattern of the depletion laser was successfully secured throughout the entire volume of the sample, including the top and bottom.
- This study presents the first-ever full volume 3D image of the mouse oocyte, which was acquired using a 2-channel 3D STED method.

**GRAPHICAL ABSTRACT:** Introducing an extended sample preparation procedure resulted in an outstanding optical quality sample environment pronounced by high refractive index, high transparency, and minimal spherical aberration. This procedure allowed 3D STED within the entire oocyte.

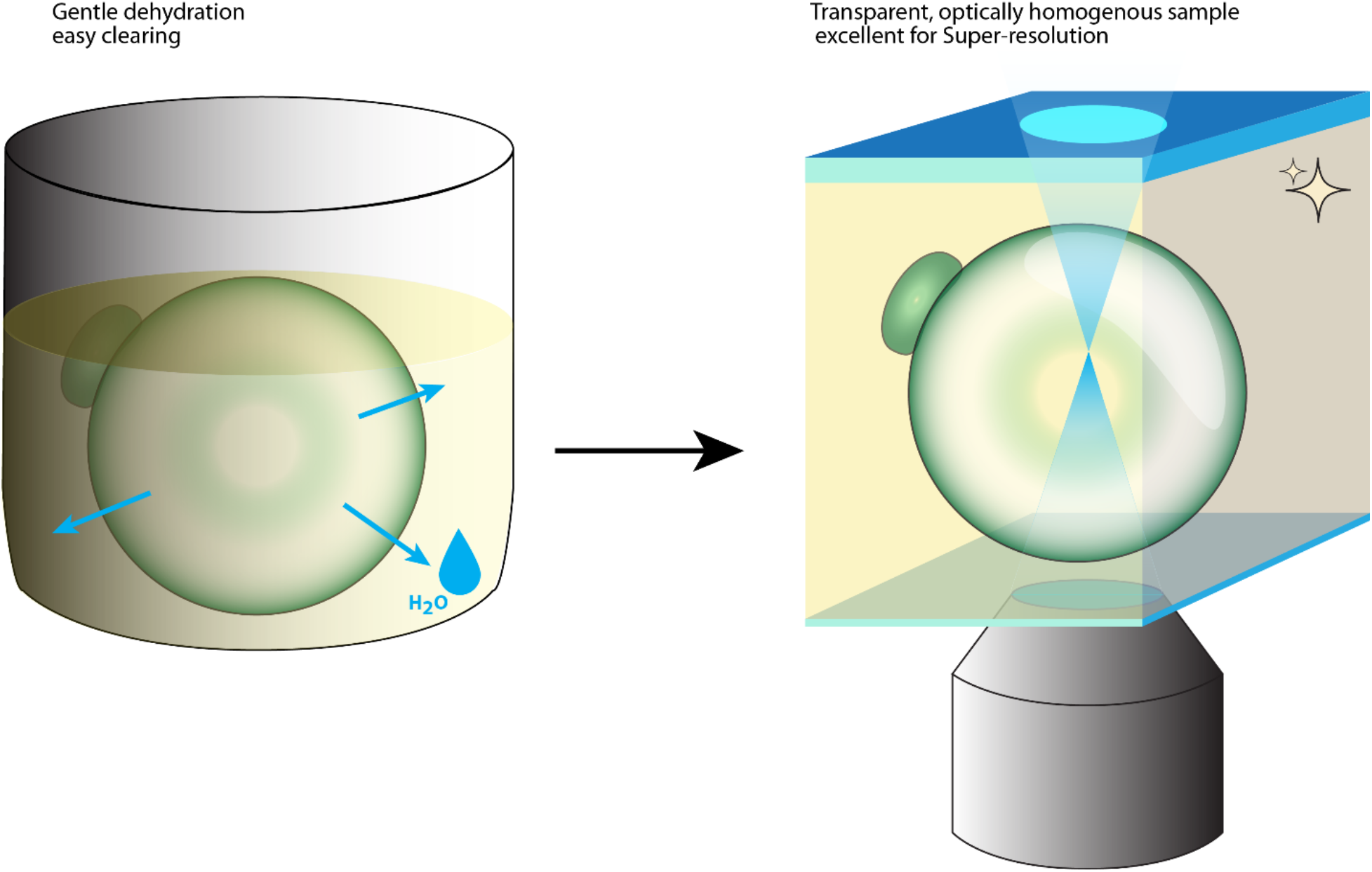

## INTRODUCTION

For many reasons mostly based on a motivation to study the context of biological structures in high detail, there is a desire for high-resolution and super-resolution (SR) imaging methods in the science. In the field of light microscopy, there are a variety of super-resolution (SR) methods based on several main approaches which allow us to move beyond the diffraction limit barrier (Schermelleh, Heintzmann et al. 2010). The trend in biology is to visualize more than the adherent cell monolayer and thus image larger and thicker samples with the advantage of a relatively deep optical sectioning in the Z-axis and confocal high-resolution (Lam, Cladiere et al. 2017). However, high-resolution confocal imaging cannot provide an image with real SR detail in nanoscale. Stimulated emission depletion (STED) microscopy is based on the confocal scanning method. STED microscopy uses a confocal scanning microscope equipped with an additional depletion laser to achieve a resolution beyond the diffraction limit (Hell and Wichmann 1994, Klar, Engel et al. 2001). The doughnut-shaped depletion laser is carefully co-aligned with the excitation laser and in the overlapping region, stimulated emission depletion of fluorochromes occurs. This phenomenon effectively shrinks the region of sample excitation and thus increases the final lateral resolution of the acquired images (Klar, Jakobs et al. 2000, Hell 2009). Moreover, higher axial resolution may be achieved by using a 3D STED technique. In 3D STED, a phase plate is inserted into the depletion laser beam path, which leads to the introduction of a so called z-doughnut shape to the imaged volume, which leads to the enhancement of axial resolution of the final image (Hein, Willig et al. 2008).

The biological scope of this research was to examine at nanoscale resolution the relationship of two key oocyte surface proteins, Juno (Bianchi, Doe et al. 2014) and CD9 (Kaji, Oda et al. 2000, Le Naour, Rubinstein et al. 2000, Miyado, Yamada et al. 2000). As we cannot solve the inherent problems with the classic confocal imaging method described above, we decided to experiment using the 3D STED super-resolution (SR) microscopy technique, which had never been used to image an oocyte in its entire volume. The main limitation of designing an experiment in oocyte imaging is sample preparation.

The oocyte is the largest cell in the mammalian body. It is perfectly round, full of cytoplasm and has an average diameter of 80 μm. The oocyte plasma membrane, called the *oolemma,* is exposed to high pressure by the contents of the oocyte and this makes oocytes fragile and sensitive to manipulation. *In vivo,* the resistance of the oolemma to cytoplasmic pressure is supported by an extracellular envelope called the *zona pellucida* (Brayboy and Wessel 2016). During the procedure of immunostaining the *zona pellucida* might have to be removed to prevent non-specific binding of antibodies and this makes oocytes extremely fragile. The critical part of the oocyte immunostaining procedure is the final transfer of an oocyte from the water-based staining solution to a high refractive index mounting medium. In adherent cells, a common procedure for this transfer is drying the sample, which ensures a homogeneous penetration of an imaging medium into the sample avoiding potential refractive index mismatches and downstream optical aberrations (Egner, Schrader et al. 1998). However, the process of drying an oocyte results in the collapse of its fragile structure. Therefore, in order to prevent oocytes prepared in this way from collapsing, it is necessary to perform the transfer of the oocyte from the water-based staining solution to the mounting media through a series of glycerol gradient solutions (Flemr and Svoboda 2011). This commonly used procedure, we call *Standard protocol* in this text. It became clear that even using this more complex procedure, the insufficient removal of water from the body of the oocyte and its surroundings could result in a significant deterioration in the quality of microscopic images obtained from the deeper parts of the sample. The oocyte is a very complicated biological specimen for sample preparation. SR techniques require high optical quality of the sample. Therefore, the combination of oocyte sample preparation and SR technique is a challenging combination. The 3D STED technique had never been used for the imaging of an entire oocyte with surface mapping, through the full depth of the sample that is around 80 μm. The STED technique is based on confocal techniques and it one of the most robust amongst the family of SR techniques, 3D STED is very sensitive to the effect of spherical aberration. Any refractive index mismatches in the sample volume introduce artifacts (Egner, Schrader et al. 1998), which are critical for the STED excitation and depletion lasers coalignment in the imaged spot. Preventing refractive index mismatches and maintaining a homogenous optical environment throughout the entire oocyte volume has thus become a major technical challenge in our efforts to achieve the highest possible image quality. Thorough sample dehydration is critical for ensuring the best possible optical conditions for imaging of a large object like an oocyte throughout its entire volume using a high refractive index mounting medium.

One way to achieve sample dehydration is through a series of solutions of increasing alcohol concentrations. The procedures based on alcohol dehydration are well known for sample preparation of germ cells (including oocytes) for electron microscopy (Schatten 2004, Nagy, Varghese et al. 2019) and fluorescence in situ hybridization (Sarrate and Anton 2009). The ethanol or methanol dehydration steps in these protocols do not introduce any structural deformations to the sample. However, the alcohol dehydration step is not a common technique used in typical oocyte immunostaining procedures for light microscopy.

In this paper, we present the successful implementation of novel steps for sample preparation that lead to the high optical quality of an oocyte sample suitable for super-resolution techniques. Even though an oocyte was used in these experiments, we believe that these sample preparation steps used for SR have a high potential for use in a wide variety of biological specimens.

## RESULTS

To address the technical difficulties in sample preparation of an oocyte for 3D STED microscopy, we had to overcome two main challenges. Firstly, it was necessary to remove water from the sample without air drying. This is because the air-drying technique always leads to the collapse of the oocyte structure, and secondly, to transfer the oocyte to a high refractive index mounting medium.

### Extended protocol implementation

We established an extended sample preparation technique that we named – *extended protocol*, based on the common fixation method and consequential steps of immuno-staining and washing. Instead of direct mounting of an oocyte surrounded and filled with water residua, we introduced gentle alcohol dehydration of the specimen with 50% absolute ethanol, followed by gradual transfer through 50% ethanol/50% 2,2’-Thiodiethanol (TDE) and 100% TDE. The sample was then added to the mounting medium AD-MOUNT C (ADVI, Ricany, CZ) which has a refractive index resembling glass (RI 1.518). The procedure is summarized in Figure 1b. The *standard protocol* (Flemr and Svoboda 2011) showing glycerol gradient transfer is shown in Figure 1a.

**Figure 1:**
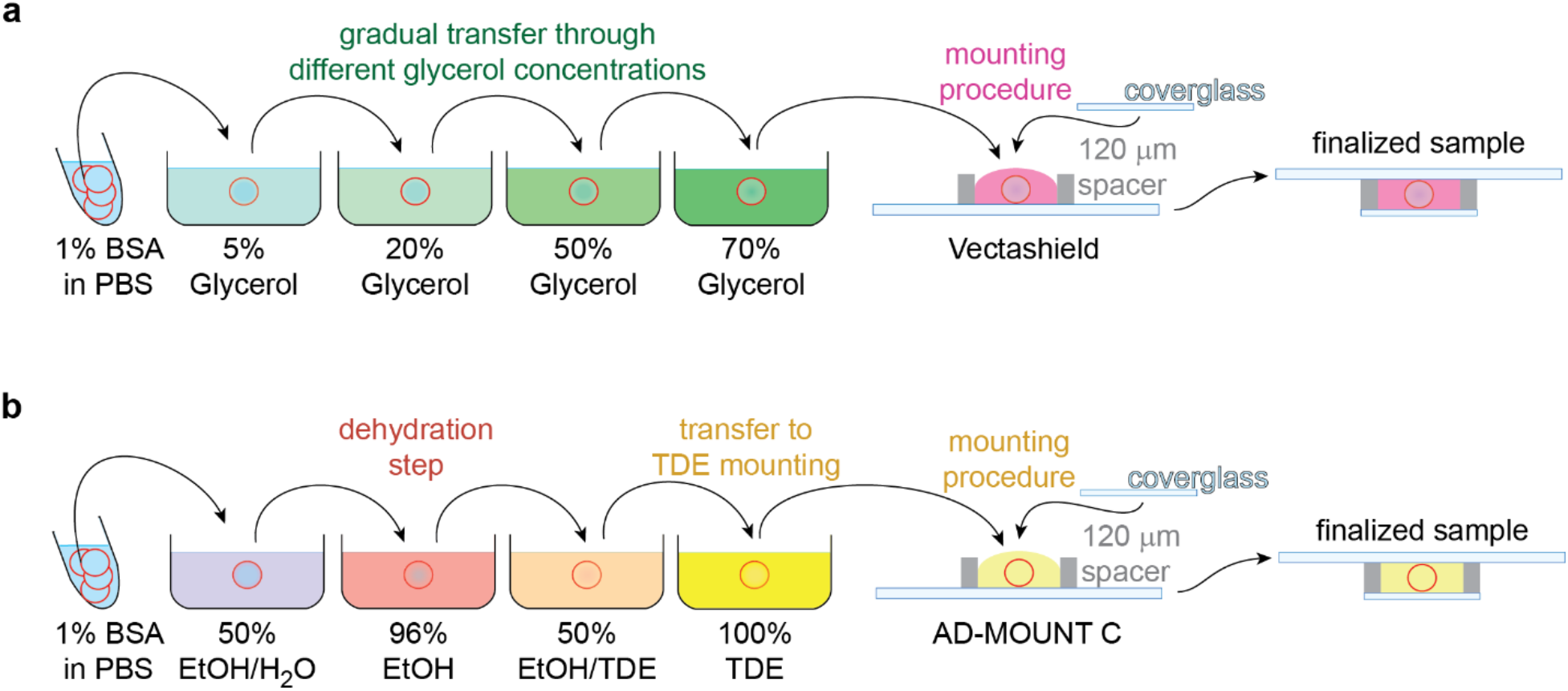
The scheme visualizes the final mounting workflow procedure within the *Standard protocol* and *Extended protocol*. **a:** *Standard protocol,* the transfer of the sample (red circle) from the water-based buffer through glycerol solutions from 5% to 70% with 10 min incubation in each step to equilibrate the gradient between the glycerol solution and sample surrounding environment. Finally, the sample is mounted to Vectashield mounting medium (Vector Laboratories, Burlingame, CA, USA). **b:** *Extended protocol*, the sample is transferred from the water-based buffer through 50% ethanol to the 96% ethanol solution for dehydration. The sample is then transferred from the ethanol through the 50% TDE/50% ethanol to the 100% TDE. In each step, the sample is incubated for 20min. Finally, the sample is mounted to the AD-MOUNT C mounting medium. After the procedure, the sample is totally clear and without refractive index mismatch gradients.

### Extended protocol preserves the signal intensities within entire sample

To verify the benefits of the *extended protocol* for oocyte SR imaging, we decided to compare optical properties of a samples prepared using the *standard protocol* and the *extended protocol*. Technically, for oocyte surface visualization, we indirectly immunolabeled the Juno protein (Bianchi, Doe et al. 2014) on the oocyte surface and divided oocytes into two experimental groups. One group of oocytes was prepared according to the *standard protocol* and the other group according to the *extended protocol*. For visualization of the oocyte groups, we used a confocal scanning microscope. After the acquisition of the oocyte images, we inspected the image quality within the entire sample volume. As a reference to optical homogeneity and transparency, we compared the signal intensities of acquired structures on the z-axis profile of the point spread function (PSF) shape on the opposite sides of the oocytes. Normally, the optical quality decreases along with the depth of the sample. When using a microscope to compare images of an oocyte, precise definitions of the cell’s different components are crucial. To facilitate accurate interpretation of microscopy data, we refer to two sides of the oocyte as ‘*top*’ and ‘*bottom*’. The term ‘*bottom*’ is used to describe the part of the oocyte that is closest to the microscope objective and the coverslip. The term ‘*top*’ is used to describe the opposite side of the oocyte, which is furthest away from the coverslip. These terms are solely used to describe the orientation of oocyte with respect to the microscope objective and do not relate to any intrinsic properties of the oocyte.

The oocytes from both groups were visualized using a confocal scanning microscope to obtain a z-stack image of the entire oocyte volume. The maximum projection and XY and ZY (orthogonal) sections of acquired Z-stacks are shown in Figures 2a, b. The oocytes prepared according to the *standard protocol* (Fig 1a) gave the expected results showing sharp PSF in the *bottom* part, while in the deeper *top* part of the specimen there was apparent loss of signal intensity and resolution and intensive blurring. In contrast, the second group of samples prepared according to the *extended protocol* (Fig 1b) showed an almost uniform shape of the PSF in *bottom* and *top* parts of the oocyte. The signal intensities were also preserved; the contrast of the image was constant throughout the entire imaged volume. The comparison of the *extended protocol* with the *standard protocol* resulted in the significantly better optical quality of the sample in respect to signal intensities, contrast, blurring, and PSF shape in the group of oocytes prepared using the *extended protocol*.

**Figure 2:**
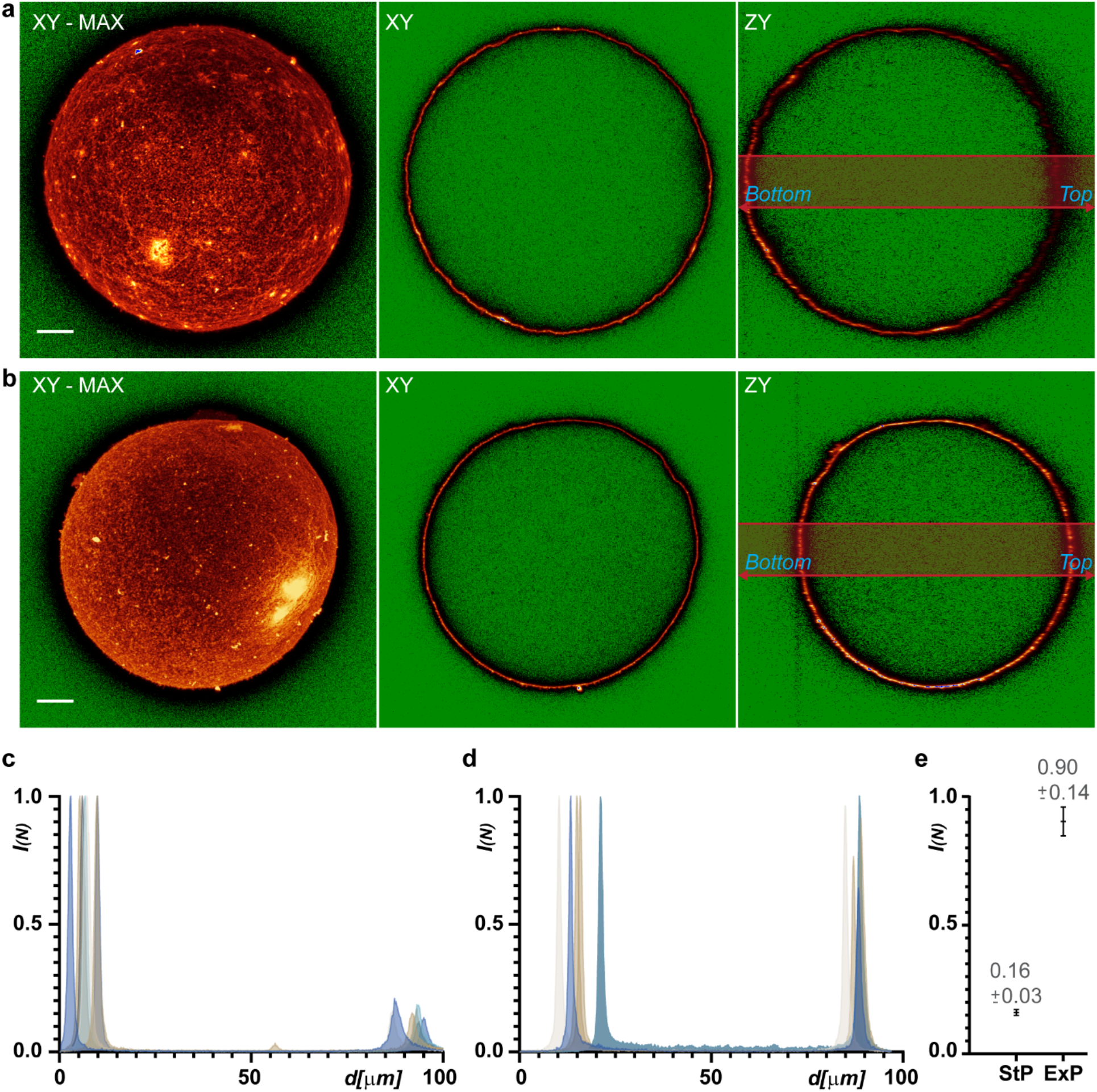
The comparison of differences in signal intensity and its fading in deeper layers demonstrated on confocal images of oocytes prepared according to the *standard protocol* and *extended protocol*. The deeper part of the object means the layers situated at regions far from the objective position, right part of the ZY orthogonal views, marked as *top*. The regions of the oocyte which is close to the objective is marked as *bottom.* **a, b:** representative confocal sections in XY and ZY (orthogonal) layers and XY – MAXIMAL projection images of the oocytes prepared according to the (**a**) *standard protocol* and according to the (**b**) innovative *extended protocol.* The lookup table “glow” (ImageJ standard) was used to more intuitively visualize the changes in signal intensities: the green color indicates the zero values, the colors from dark red, red, through yellow up to white cover the dynamic range of the image, and the maximal values (image saturation) are highlighted by the blue color. The bar represents 10 μm. The *standard protocol* (**a**) shows the effect of spherical aberration propagated in signal fading and smearing of structures in *top* of the object. The innovative *extended protocol* (**b**) by comparison demonstrates minimal spherical aberration and the clearing effect results in preservation of signal intensities with almost no visible fading effect and a fairly homogenous shape of structures in the *top* of the specimen. **c, d:** charts visualize the normalized signal intensity *“I_(N)_”* on Y axis against the values on X axis where the distance *“d”* into the sample represents the plot profiles in ZY orthogonal views of confocal images. The chart (**c**) represents the plot in the *standard protocol* and (**d**) in the innovative *extended protocol.* (**a, b**) The signal intensities represent area under the light red-gray rectangle in the YZ orthogonal projections. Area measurements were preferred over simple line profiles to minimize the noise of the plotted data. **e:** the chart represents the average decrease of intensities *“I_(N)_”* on *top* area normalized to the intensities on *bottom* area. The average and standard deviation is plotted, and the corresponding numbers visualized for oocytes processed by the *standard protocol* (StP) and the *extended protocol* (ExP).

To better illustrate the differences in fluorescence signal intensity between the *bottom* and *top* parts of the oocytes prepared according the two protocols, the signal intensities under selected rectangular areas (Fig. 2a, b) were quantified and plotted to graphs, see Figures 2c and d. These graphs show the signal intensities from the *bottom* part, on the picture situated on the left side, to the *top* part, on the picture situated on the right side of the oocyte orthogonal image. The plotted signal intensity values were calculated from the YZ orthogonal projections of volume images.

The signal intensity profile of the sample prepared by the *standard protocol* showed a significant attenuation of the fluorescence signal in the deeper layers of the sample (Fig. 2c), while the signal intensity remained almost constant in the sample prepared using the *extended protocol* (Fig. 2d). A quantitative comparing the normalized signal intensities from the *bottom* to *top* were plotted (Fig. 2e) and this illustrates the preservation of the fluorescent signal throughout the oocyte volume when the *extended protocol* was used.

### Extended protocol ensures the lasers PSF co-alignment within the sample

The key prerequisites for successful 3D STED imaging of a large biological sample (in this case an oocyte) are its high transparency and the relatively constant shape of the PSF throughout its volume. These conditions ensure the correct formation of both the XY doughnut and the Z-doughnut, the precise co-alignment of the depletion laser doughnuts with excitation PSF, and preservation of signal intensity in the deeper parts of the sample. Inspired by the previously cited approach for visualization of 3D STED alignment in a volume sample, we have prepared the most realistic calibration sample with capability to visualize both excitation and depletion lasers PSFs. To visualize the oocyte surface, we immunostained the Juno protein (AlexaFluor 488) and in the next step after the staining procedure we adhered 80 nm gold beads directly to the oocyte surface. Then we processed the oocyte sample using the *extended protocol.* Oocyte visualization was performed in standard confocal fluorescence mode and adhered gold beads were visualized in confocal reflection mode (Fig.3a). The results showed that the PSF shapes of a single bead located both on the *bottom* or on the *top* are very similar, both for the excitation and depletion laser and in 2D and also in 3D STED mode (Fig.3b). This means, that the sensitive co-alignment of excitation and de Nynäshamnpletion lasers are conserved within the full specimen volume. This suggests the 3D STED imaging capability within the full depth of the sample prepared using the *extended protocol*.

**Figure 3:**
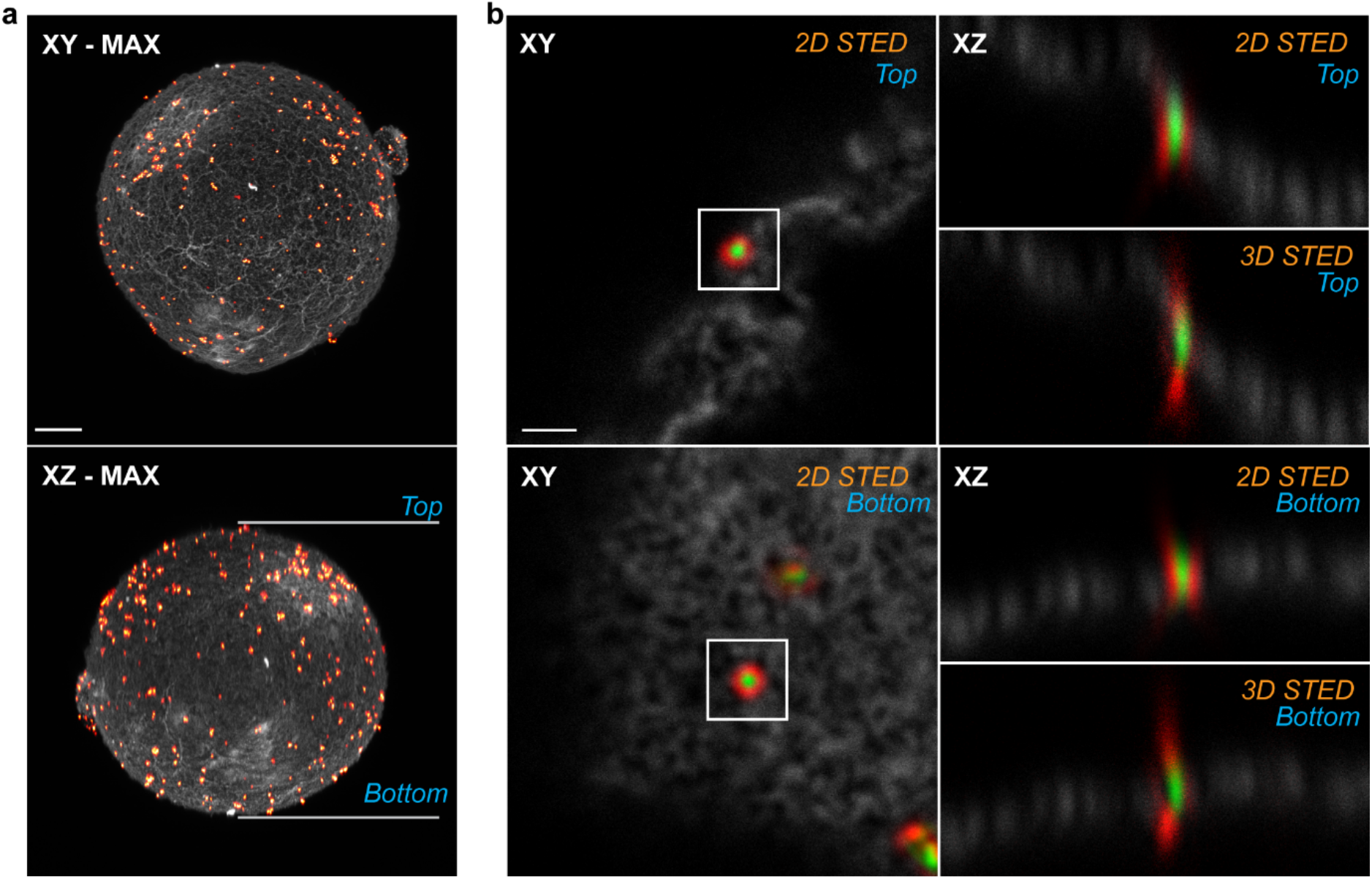
Confocal images visualize the oocyte with adhered 80 nm gold beads. The oocyte sample was prepared using the *extended protocol.* The image shows the appropriate formation of 2D STED or 3D STED depletion laser pattern in the area from a few μm to approx. 80 μm in depth. **a:** maximum intensity projection of the oocyte confocal images in XY and XZ axis. Grey channel represents fluorescence signal of the counterstaining for visualization the oocyte, the “Red Hot” lookup table represents the visualization of adhered 80 nm gold beads acquired in reflection. The bar represents 10 μm. **b:** confocal sections visualize the membrane of the oocyte in the gray channel. Gold beads on the membrane were used for visualization of the PSFs of excitation laser (green) and depletion laser (red). The XY confocal section shows the lateral co-alignment and the XZ confocal scan shows the axial coalignment of excitation and depletion laser. The position relative to the objective is marked as *bottom* or *top*. The marks “2D STED” or “3D STED” indicates the settings of the microscope super-resolution STED module in the images with the corresponding depletion laser PSF shape formed on gold beads. The bar represents 1 μm.

### 3D STED applicable within entire specimen after extended protocol sample preparation

To verify the applicability of the *extended protocol* for 3D STED imaging, we used oocytes with immunolabeled membrane proteins Juno (Bianchi, Doe et al. 2014) and CD9 (Kaji, Oda et al. 2000, Le Naour, Rubinstein et al. 2000, Miyado, Yamada et al. 2000), which important for our research interest. Immunolabeled oocytes were processed by both the *standard* and *extended protocols*. A comparison of standard confocal images and super-resolution 3D STED images obtained by application of 60% z-doughnut is shown in Figure 4. The application of 3D STED significantly increased both lateral and axial resolution in the *bottom* of the specimen, regardless of the protocol used, while significant differences are apparent in the *top* of the specimen (Fig 4a, b). Specimens prepared using the *standard protocol* exhibit a dramatic loss of signal intensity combined with smearing of the signal localization on structures on the *top* see Figure 4a of the specimen; this procedure of sample preparation is not effective for imaging of the entire oocyte by 3D STED technique. In contrast, oocytes processed using the *extended protocol* show excellent preservation of signal intensity and resolution in both the *bottom* and the *top* regions; structures are resolved in a comparable manner throughout the entire oocyte volume, see Figure 4b. These results demonstrate that the *extended protocol* is a sample preparation technique suitable for SR imaging of large biological structures; the technique allows visualization of the entire oocyte volume using 3D STED super-resolution microscopy.

**Figure 4:**
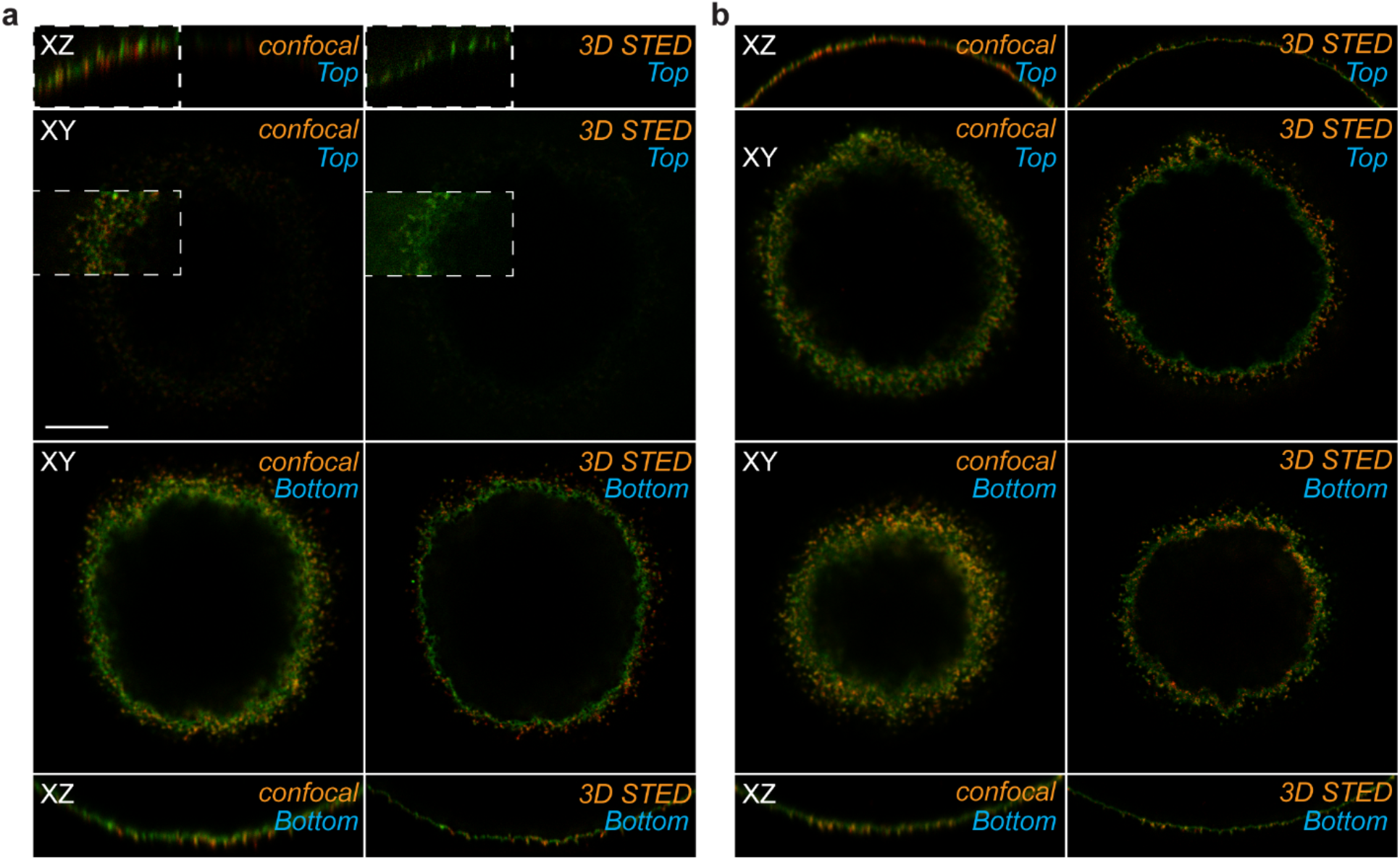
Confocal and 3D STED images of the oocyte prepared using the *Standard protocol* and the *extended protocol*. The image represents the simple XZ orthogonal scan on the first row, the second row shows a single XY section, both in the *top*. The third row of the image visualizes a single XY section in the position *bottom* of the oocyte, the bottom row of the image shows the simple XZ orthogonal scan of the part *bottom.* The method used for the acquisition is named “confocal” or “3D STED”. The bar represents 10 μm. **a:** *Standard protocol* gives images with standard confocal quality in the *bottom* part, which is also suitable for 3D STED super-resolution imaging. In contrast, on the *top* side, the signal intensity is poor and fades away; the 3D STED super-resolution imaging on that side failed. To only improve the visualization and demonstrate the weakness of the signals, we non-systematically increased the intensities of the images in the white-bordered squares which are on the *top* side. **b:** *Extended protocol* results in standard confocal and 3D STED super-resolution images with fully comparable quality on both *bottom* and *top* parts, without any major fading of signal intensities.

### Comparable resolution within entire specimen after extended protocol sample preparation

Our next question was how the resolution differs between the *bottom* and *top* of the specimen. For the quantification of the resolution, we used the Fourier ring correlation (FRC) computation method (Nieuwenhuizen, Lidke et al. 2013). For the calculation, we acquired optical sections in both the *bottom* and *top* regions. The FRC measurement was performed on raw data. For visualization, the images were conservatively deconvolved to reduce the unwanted noise (Fig.5a). To ensure robustness of the resolution measurement, the data from five independently prepared and acquired oocyte images were used. The results of the experiment showed that the image resolution acquired by STED microscopy in both regions, *bottom* (70.6 ±8.0 nm) and *top* (76.6 ±9.2 nm), was comparable.

**Figure 5:**
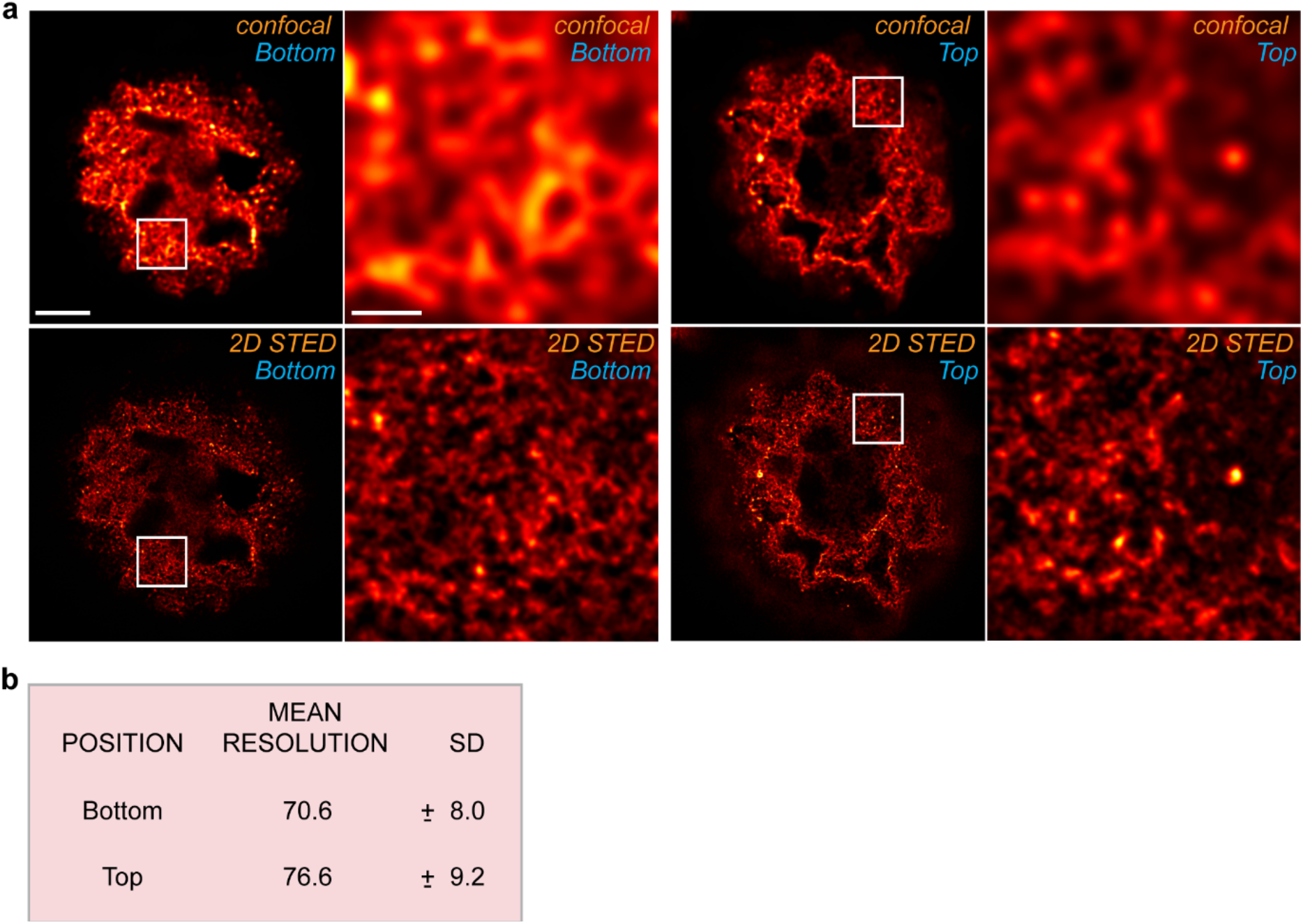
The image resolution achieved with 2D STED approach within the oocyte sample. **a**: the comparison between the confocal (marked with *“confocal”)* and 2D STED (marked with “*2D STED*”) acquisition method in *bottom* and *top* positions of the acquired oocyte. The bar represents 10 μm at overviewed image and 1 μm at the zoomed area. **b**: the table quantifies mean and standard deviation of the resolution calculated by FRC method within the image acquired in part *bottom* or *top* of the specimen.

### 3D STED acquisition of entire oocyte allowed by extended protocol sample preparation

For the complete acquisition by two-channel 3D STED microscopy technique, the oocyte sample was prepared according to the *extended protocol.* After the acquisition, the raw images were deconvolved using the STED option of Huygens Professional software. The deconvolution provides the data with maximal recovered acquired structures, improves signal to noise ratio, and restores SR image quality. The final images clearly show the details in the distribution of both proteins, Juno and CD9, over the entire oocyte surface. The technique allows the signal localization at nanoscale resolution and with constant high resolution through the entire specimen (Fig.6a and Supplementary video1). The final super-resolution images reveal extremely fine microvillar structures (Fig.6b) that have so far only been visualized by electron microscopy (Benammar, Ziyyat et al. 2017). The individual oolemma compartments, such as the microvillar membrane and planar membrane located between individual microvilli, can clearly be distinguished and we were able to demonstrate (Frolikova et al. – manuscript in preparation) the differences in localization of these two proteins in the individual oolemma compartments (Fig.6c).

**Figure 6:**
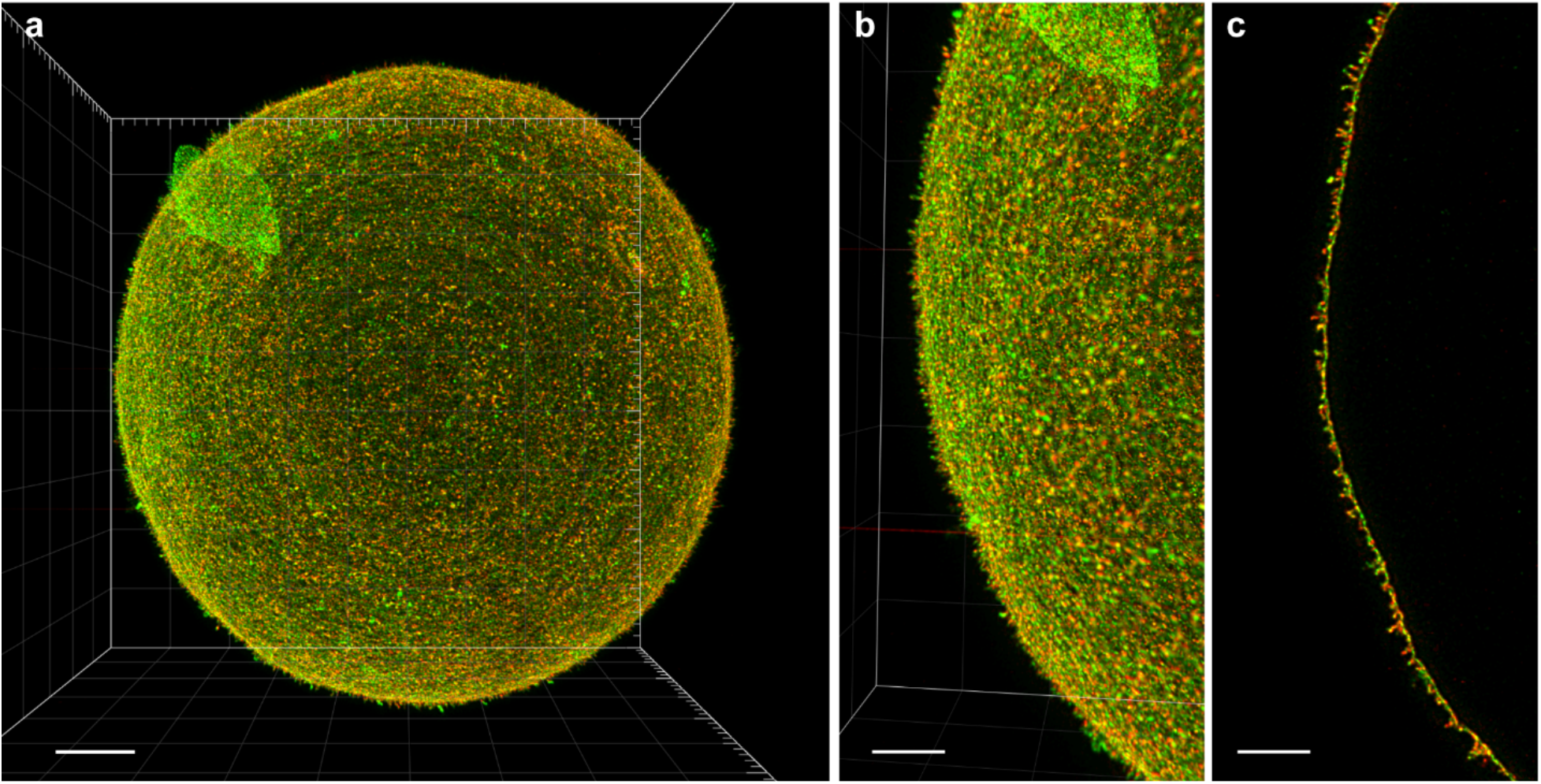
The visualization of the final deconvolved 2-channel 3D STED super-resolution volume reconstructed image of the entire mouse oocyte. The image shows an overview of the oocyte and reveals the detail of microvillar structure visualized with immunoflurescent staining of Juno (green) and co-localized CD9 protein (red) in the zoomed area. **a, b:** (**a**) 3D maximal projection of the final full z-stack of the entire oocyte in 3D STED microscopy and (**b**) close-up of the selected part of entire oocyte image. The bar represents (**a**) 10 μm and (**b**) 5 μm. **c:** a representative maximal intensity projection of 10 selected optical sections around the equatorial layer of the oocyte in super-resolution 3D STED image. The bar represents 5 μm.

In this paper, we present the volume 2-channel 3D STED image of the mouse oocyte. The detailed visualization of Juno and CD9 proteins shows their co-distribution at nanoscale resolution within the microvillar surface structures all-over the entire oocyte. Altogether, the results provide evidence that the *extended protocol* is a fully suitable sample preparation procedure using the SR microscopy technique – 3D STED for imaging the oocyte, which is a large and extremely fragile specimen.

## DISCUSSION

The high-resolution and super-resolution (SR) techniques in light microscopy reveal very fine biological structures, their compositions and allow studying of co-localization to a nanoscale dimension. SR methods are suitable for many types of specimen, nevertheless, the limitation of the methods is the thickness of the sample and depth of the region to be imaged within that sample. The confocal based method STED naturally removes the out-of-focus light and thus allows the acquisition of deeper layers of the sample (Valli, Garcia-Burgos et al. 2021). While the 3D STED method improves the resolution not only in lateral but also in an axial orientation (Osseforth, Moffitt et al. 2014), the method is also limited by sample thickness; for the 3D STED method the overlapping of the excitation and depletion laser PSF pattern is crucial. In deeper layers of the sample, the geometry of the excitation and depletion beams are differently affected and it impairs the method (Jacquemet, Carisey et al. 2020). Keeping in mind that the limitation here is mostly caused by refractive index mismatch and down-streamed spherical aberration, the crucial achievement what we achieved in our approach is the method of sample preparation. The procedure we named here as *extended protocol* where we focused on the two steps essential for resulting in optical homogeneity of the final sample, first, the removal of water residues, second, appropriate mounting.

The removing of the water from the sample according to the previous standard procedure for oocyte sample preparation for fluorescence microscopy is based on a gradual exchange of water from the sample through several steps with increasing glycerol concentration and the final assembly of the sample into the Vectashield mounting medium (or qualitatively similar) (Flemr and Svoboda 2011). This method is suitable for robust confocal microscopy and using some method of compensation of the light attenuation with depth of images (Capek, Janacek et al. 2006). Anyway, the method is not sufficient for the 3D STED method which is more sensitive, and the later compensation protocol does not work while the principle of the method is affected by spherical aberration. In contrast, our *extended protocol* very efficiently removes residual water from the sample by means of alcohol-based dehydration. During the procedure the specimen never dries out and as a result, the fragile structure of the oocyte does not collapse.

The second key step in the *extended protocol* is the mounting of the sample into the AD-MOUNT C mounting medium with a glass-like (RI), 1.518. This mounting medium minimizes the effect of spherical aberration and results in an almost uniform PSF shape throughout the depth of the sample (in the case of an oocyte specimen, around 80 μm). The fact that the RI equals glass, is highly important for the method; even though the sample is optically homogenous and clear, in case the mounting medium RI does not match glasses RI, it is necessary to compensate for the effect of spherical aberration using the objective correction collar to ensure correct PSF shape in deep areas of the sample and 3D STED in-depth alignment (van der Wee, Fokkema et al. 2021). With the AD-MOUNT C, TDE-containing mounting medium, the PSF and thus the 3D STED alignment stays uniform within the whole volume of the sample. Additionally, TDE has a rapid clearing effect (Aoyagi, Kawakami et al. 2015). This minimizes the scattering of light when passing through a thick sample making the biological structures more transparent. For excitation, depletion, and emission light, the fluorescent signal is uniformly preserved throughout the depth of the sample.

The water removal significantly improves the optical quality of the sample, while the dehydration process without drying ensures the integrity of the sample structures and preserves the shape properties of the object. The mounting of the sample into a high refractive index mounting medium with the clearing effect leads to an optically homogenous and transparent sample environment. This makes the technique suitable for high-resolution or super-resolution microscopy methods. We succeeded with 2-channel 3D STED imaging throughout the full oocyte volume in constant resolution. This achievement may be of special interest because oocytes are a widely used model for membrane proteins, e.g., auxin hormone transporter studies reviewed in 2018 (Barbosa, Hammes et al. 2018). Generally, in science, the trend is to convert from 2D adherent cell culture to more realistic models like organoids (Kim, Koo et al. 2020). We believe that this simple technique would have a potential for a much wider range of applications in the fluorescence microscopy field, including other super-resolution techniques and other types of thicker biological specimens.

## MATERIAL AND METHODS

### Animals

Inbred C57BL/6J female mice were housed in a breeding colony of the Laboratory of Reproduction, IMG animal facilities, Institute of Molecular Genetics of Czech Academy of Science. Food and water were supplied ad libitum. The female mice used for all experiments were healthy, 23–26 days old, with no sign of stress or discomfort. All animal procedures and experimental protocols were approved by the Animal Welfare Committee of the Czech Academy of Sciences (Animal Ethics Number 66866/2015-MZE-17214, 18 December 2015).

### Oocyte preparation

Female mice were hormonally stimulated with 5UI PMSG—Pregnant Mare’s Serum Gonadotropin (Folligon, Intervet International B.V., Boxmeer, The Netherlands) at 15:00 (the eighth hour of light cycle) on the first day of the protocol. 5UI of hCG—human Chorionic Gonadotropin (CG10, Sigma-Aldrich, St. Louis, MI, USA) were applied to mice at 1pm on the third day of the protocol (46th hour after using PGMS). After 12 hr., the female mice started ovulating. At 9am on the fourth day of the protocol, female mice were sacrificed by cervical dislocation and both ampullas of fallopian tube were isolated and placed in preheated M2 medium (M7167, Sigma-Aldrich^®^, St. Louis, MI, USA). Cumulus-oocytes complex (COC) was released into the M2 medium by ampulla tearing. In the next step, for releasing cumulus cells, COC was transferred into a fresh 100 μl drop of M2 medium with hyaluronidase (concentration 0,1 mg /ml) (hyase, from bovine testes, H4272, Sigma-Aldrich^®^), covered with high viscous paraffin oil (P14501, Carl Roth, Germany) and left in the incubator (set on 37 °C, 5% CO_2_) for 10 min.

After washing, the cumulus free eggs were transferred into a drop of Tyrode’s solution (T1788 Sigma-Aldrich^®^) to remove *zona pellucida.* The zona free eggs were washed 2× in 1 % BSA in PBS and fixed with 3.7 % paraformaldehyde (P6184, Sigma-Aldrich^®^) for 20 min and washed 2× in 1% BSA in PBS. The oocytes were incubated overnight in 4 °C in a drop of primary antibodies rat monoclonal anti-CD9 (KMC8.8) (sc18869, Santa Cruz Biotechnology, Inc., Dallas, TX, USA) diluted 1:50 in 1% BSA in PBS and rabbit polyclonal anti-Folate receptor 4 (Juno) (abx102438, Abbexa, UK) diluted 1:50 in 1% BSA in PBS followed by 1 hr. incubation with secondary antibodies anti-rat IgG Abberior STAR 635P (Abberior GmbH, Germany) and anti-rabbit IgG Abberior STAR 580 diluted 1:100 in 1% BSA in PBS at room temperature (RT) and washed 2× in 1% BSA in PBS.

### Adhesion of 80 nm gold beads to the oocyte surface

Gold beads with a volume of 500 μl of stock solution (Gold nanoparticles, 80 nm diameter, OD 1, stabilized suspension in 0.1 mM PBS, 753661 Sigma-Aldrich^®^) were pelleted (RT, 10 000 *g*) and re-suspended in 500 μl 1× PBS buffer. Immunostained oocytes were transferred into the gold beads suspension before the mounting procedure. The oocytes were incubated 20 min at RT with gentle agitation. After the incubation, the oocytes were washed 3× in PBS and processed according to the *extended protocol* mounting procedure.

### Oocyte mounting procedure: standard protocol

The samples were mounted according to the standard procedure for the light microscopy (Flemr and Svoboda 2011); the oocyte was transferred from the 1% BSA in PBS used for washing through a series of 5%, 20%, 50%, and 70% glycerol solutions, incubated for a minimum of 10 min at each step and finally mounted to the commercially available mounting medium Vectashield (Vector Lab., Burlingame, CA, USA).

### Oocyte mounting procedure: extended protocol

In this procedure, gradual ethanol (EtOH) dehydration steps and gradual transfer to the mounting medium were added before final specimen mounting; the oocyte stored in 1% BSA in PBS was transferred to the 50% EtOH in water, incubated for 10 min at RT, then the oocyte was incubated in 96% EtOH for 20 min at RT. From the ethanol, the oocyte was transferred to the 50% TDE in EtOH, incubated for 20 min at RT, and finally transferred to the 100% TDE and incubated for 20 min at RT. The oocyte was transferred to the cover glass and then mounted to the TDE based mounting medium AD-MOUNT C (ADVI, Ricany, CZ). For observation and handling of oocytes throughout the procedure, a standard stereomicroscope was used in the initial steps. Nevertheless, because of the dehydration and transfer of oocytes to solutions with high nD in the final steps of the procedure, oocytes started to be nearly invisible in standard bright field/dark field or opaque contrast. Therefore, use of the fluorescence imaging mixed with the opaque contrast was necessary for visualization and handling of oocytes in the final steps of the procedure.

### Confocal scanning and super-resolution acquisition

For both classical confocal scanning and STED super-resolution acquisition, the Leica TSC SP8 STED 3× microscope equipped with pulse white light laser (WLL2) for the excitation and pulse 775 nm laser for the emission depletion was used. Image acquisition was performed using the software LAS X 3.5.6 (Leica Microsystems, DE).

Acquisition settings for confocal imaging at Fig.2: 100× 1.4 NA STED oil objective, oil n = 1.518, pinhole 1AU according to the excitation wavelength 580 nm, pixel size XY: 175 nm × 175 nm with z-step size 250 nm, detection on Hybrid Detector (HyD) in photon counting mode, emission captured in the interval 587-617 nm, interval for the time gating window: 0.3-10 ns.

For the Figure 3, the confocal and 3D STED comparison was needed. The same pixel size was used for standard confocal and SR imaging. For the XZ orthogonal view acquisition, the XZ scan feature of the microscope was used. Confocal settings: 100× 1.4 NA STED oil objective, oil n = 1.518, pinhole 1AU according to the excitation wavelength 580-640 nm, pixel size XY or XZ: 19 nm × 19 nm, detection on HyD in photon counting mode, emission captured in the interval 587–617 nm or 647–728 nm, interval for the time gating window: 0.3–10 ns for both channels. In 3D STED acquisition, the pulsed 775 nm depletion laser with 60% 3D STED compensated with additional line accumulation of incoming photons was used. The 3D STED acquisition was performed with 0.6 AU pinhole size.

Figure 4 showing the oocyte with gold beads adhered on its surface was acquired in combination of reflection and fluorescence settings of the acquisition mode – the visualization of excitation (488 nm)/depletion (775 nm) PSF was performed on PMT detectors (600 V) while AOBS was in reflection mode while the counter-visualization of oocyte surface (Juno protein - AlexaFluor 488) was acquired in fluorescence on HyD (photon counting mode). The visualization of depletion 2D STED “donut-like” PSF was performed with 0% 3D STED while 3D STED PSF shape was acquired with 100% 3D STED option.

Figure 5 was acquired with accent on standardized comparison of highest resolution reached in on the oocyte surface on “close” and “far” from the coverslip region. proximal and distal regions. Acquisition in 2D STED imaging mode on 100× 1.4 NA STED oil objective, oil n = 1.518, pinhole 0.6AU according to the excitation 640 nm, pulsed 775 nm depletion laser with 0% 3D STED, pixel size XY: 14 nm × 14 nm, one plain image, detection on HyD in photon counting mode, emission interval 647–728 nm, interval for the time gating window: 0.3–10 ns. For the FRC resolution calculation method, there is important to obtain two images of the same layer that differ by the noise, therefore, to minimize differences between the images, we performed the acquisition in two identical between-lines sequences.

Figure 6 was acquired as a z-stack in 3D STED imaging mode with settings: 100× 1.4 NA STED oil objective, oil n = 1.518, pinhole 0.6AU according to the excitation wavelength 580 nm or 640 nm, pulsed 775 nm depletion laser with 60% 3D STED compensated with additional line accumulation of incoming photons, pixel size XY: 22 nm × 22 nm and z-step size 94 nm, detection on HyD in photon counting mode, emission interval 587–617 nm or 647–728 nm, interval for the time gating window: 0.3–10 ns for both channels. While the calculation of the resolution was performed on raw images, for the visualization we apply gentle deconvolution to reduce the noise.

### Calculation of image resolution

The computed resolution depends on the signal-to-noise ratio, spectral image content and effective optical transfer function. Thus, it is more bias-independent compared to the more commonly used selective FWHM measurement. In principle, it is more suitable for isotropic image data; the FRC method for the measurement of the lateral resolution in XY optical sections was used (Nieuwenhuizen et al.,2013), its implementation in Fiji - ImageJ open-source software platform (https://imagej.net/Fiji). The FRC plugin is available from the PTBIOP Update Site. It requires to load two images of the same scene that differ only in noise content. Subsequently, FRC curve is calculated, and a fixed threshold value of 1/7 was used in our case to estimate the resolution.

### Software for image finalization and visualization

The image sections, maximal projections and orthogonal views were prepared in Fiji - ImageJ open-source software platform (https://imagej.net/Fiji). The raw super-resolution (SR) images of the whole oocyte were processed using the STED deconvolution module in Huygens Professional software (Scientific Volume Imaging, NL; version 20.10). The SNR for 3D STED images of the oocyte was estimated around value 7 and 5 for 580 nm and 640 nm excitations channels, respectively, number of deconvolution iterations automatically stopped on threshold value 0.05; the CMLM method was used. For the volume reconstruction, 3D visualization and movie creation, Imaris software (Bitplane AG, CH; version 9.6) was used.

## AUTHOR CONTRIBUTION

M.F. prepared all the biological samples for the 3D STED super-resolution microscopy, collaborated in optimization of the *extended protocol*, 3D STED imaging and image processing, and contributed to the writing of the manuscript. M.C., M.B. and H.C. contributed to image processing and manuscript writing. V.K. contributed to the method development and manuscript writing. E.V. contributed to the experimental part of the study. M.G. contributed to manuscript writing. K.K. contributed to writing the manuscript and provided funding. O.H contributed to manuscript writing and provided funding. I.N. developed and optimized the *extended protocol* for oocyte preparation, performed the 3D STED acquisition settings and collaborated in 3D STED imaging and image processing, contributed to writing the manuscript and provided the funding.

## ACKNOWLEDGEMENTS

The work was supported from European Regional Development Fund, project “Modernization and support of research activities of the national infrastructure for biological and medical imaging Czech-BioImaging” (No. CZ.02.1.01/0.0/0.0/16_013/0001775). Microscopy experiments, the image processing included, were performed at the Light Microscopy Core Facility, Institute of Molecular Genetics of the Czech Academy of Sciences, Prague, Czech Republic, supported by MEYS (LM2018129, CZ.02.1.01/0.0/0.0/18_046/0016045) and RVO - 68378050-KAV-NPUI. BIOCEV project (CZ.1.05/1.1.00/02.0109) from the ERDF, by the Grant Agency of the Czech Republic No. GA-22-30494S (K.K.), and by the Institutional support of the Institute of Biotechnology RVO: 86652036. V.K. was supported by the ERDF project “Plants as a tool for sustainable global development” (No. CZ.02.1.01/0.0/0.0/16_019/0000827). M.G. was supported by the Grant Agency of the Czech Republic No. 21-21736S and ID Project No. LX22NPO5102 - Funded by the European Union - Next Generation EU.

